# Patching holes in the *Chlamydomonas* genome

**DOI:** 10.1101/030163

**Authors:** Frej Tulin, Frederick R. Cross

**Affiliations:** The Rockefeller University, New York

## Abstract

The *Chlamydomonas* genome has been sequenced, assembled and annotated to produce a rich resource for genetics and molecular biology in this well-studied model organism. However, the current reference genome contains ~1000 blocks of unknown sequence (‘N-islands’), which are frequently placed in introns of annotated gene models. We developed a strategy, using careful bioinformatics analysis of short-sequence cDNA and genomic DNA reads, to search for previously unknown exons hidden within such blocks, and determine the sequence and exon/intron boundaries of such exons. These methods are based on assembly and alignment completely independent of prior reference assembly or reference annotation. Our evidence indicates that ~one-quarter of the annotated intronic N-islands actually contain hidden exons. For most of these our algorithm recovers full exonic sequence with associated splice junctions and exon-adjacent intron sequence, that can be joined to the reference genome assembly and annotated transcript models. These new exons represent *de novo* sequence generally present nowhere in the assembled genome, and the added sequence can be shown in many cases to greatly improve evolutionary conservation of the predicted encoded peptides. At the same time, our results confirm the purely intronic status for a substantial majority of N-islands annotated as intronic in the reference annotated genome, increasing confidence in this valuable resource.

## Introduction

### The Chlamydomonas reference genome and annotated transcript models

The assembled Chlamydomonas reference genome is 120 Mb long, 65% GC and very repeat-rich. The assembly contains 17 chromosomes (~1-10 Mb) and a further 37 repeat-rich ‘scaffolds’ (0.1 -0.8 Mb) (Merchant et al 2007). The reference assembly, and an annotated genome including full transcript models (Blaby et al 2014) is available on the public-access Phytozome website (/http/phytozome.jgi.doe.gov); we will refer to the sequence assembly and annotations available on this website as ‘Phytozome’ or ‘reference’. About 3.5% of the nucleotides are indicated as ‘N’ (unknown sequence). These N’s are distributed in ‘N-islands’ of varying length, mostly ~100 nucleotides; most islands are annotated as being present within intronic sequence. These islands are effectively ‘placeholders’, where the length of the island is intended to reflect an estimate of the amount of missing sequence (Merchant et al 2007).

Here we present bioinformatic and experimental methods to determine whether N-islands assigned to introns are entirely intronic, or instead contain exons missing from the assembled sequence. Our methods allow determination of the sequence of these hidden exons along with flanking intronic sequence.

### Materials and Methods

Genomic sequences and transcript models (Merchant 2007, Blaby et al. 2014) were downloaded from Phytozome. Illumina RNAseq libraries were described in Tulin and Cross (2015). Illumina genomic libraries were from parental strains in the background used for the RNAseq experiments, described in Tulin and Cross (2014, 2015). Alignment of RNAseq reads to Phytozome transcript models was described (Tulin and Cross 2015). We downloaded BLAST (Altschul e al., 1990), Trinity (Grabherr et al, 2011; Haas et al. 2013) and the Bowtie2 aligner (Langmead and Salzberg 2012). The latter was used in ‘local’ mode to align genomic reads to Trinity objects. Trinity and BLAST results were analyzed with shell scripts and Perl code. Analysis of genomic reads aligned to Trinity objects was carried out with MATLAB code. All code is available on request.

### Results

#### Targeted *de novo* transcriptome assembly

It is likely that blocks of unknown sequence (‘N-islands’) frequently result from assembly difficulties from blocks of repetitive or low-complexity sequence, which are highly abundant in *Chlamydomonas*; such sequences can cause loss of adjacent high-complexity sequence from the assembly. The Phytozome reference annotation contains 789 intronic N-islands of varying length, in 744 gene models. However, there is no guarantee *a priori* that such a block of unknown sequence does *not* contain unknown exons (Figure 1A). In this case *de novo* assembly from RNAseq reads could recover the missing high-complexity sequence, even if it is hidden in low-complexity sequence in its intronic context.

**Figure 1.**
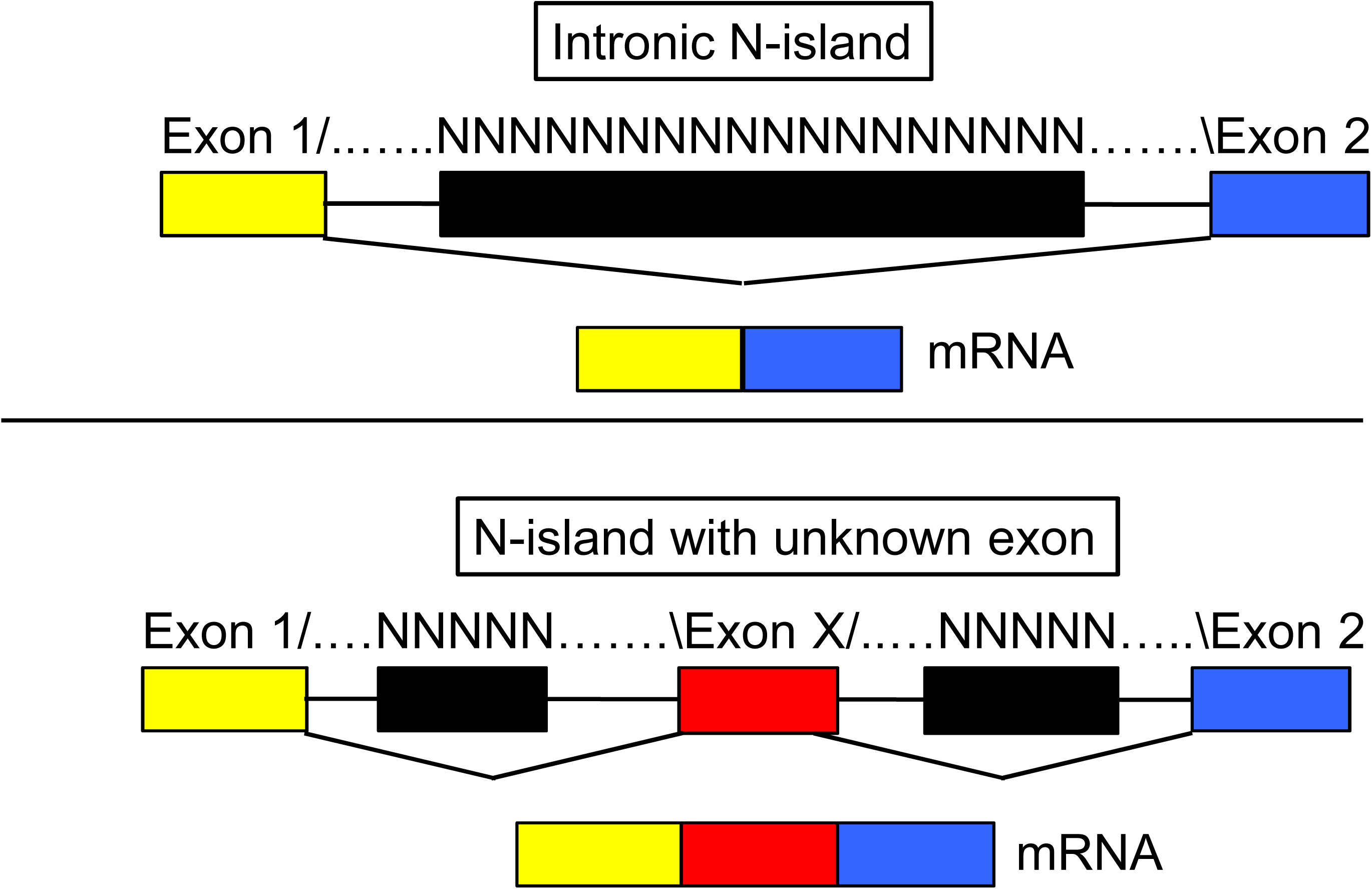
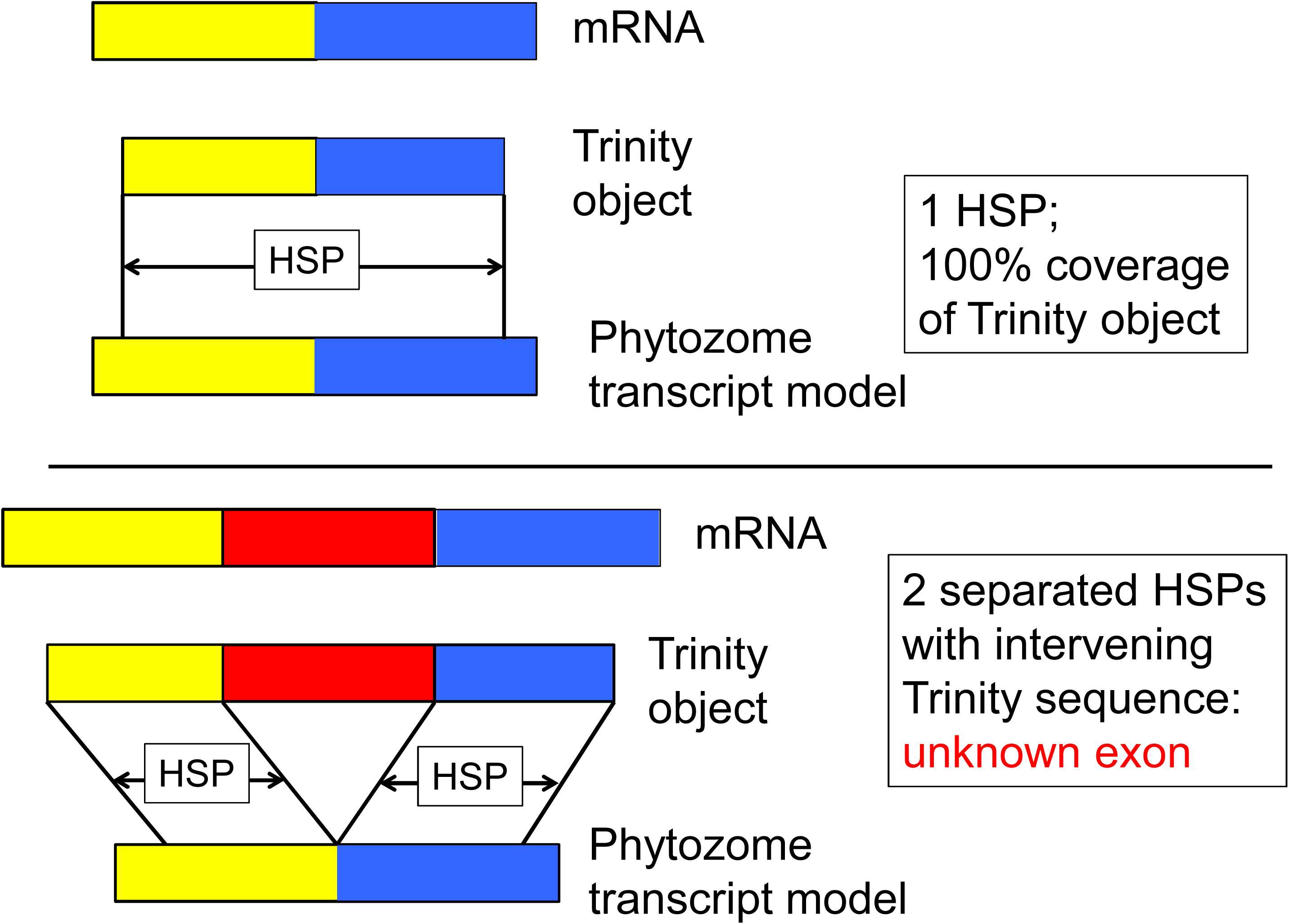
N-islands annotated as intronic can contain hidden exons: the separated HSP test. **A.** Top: Gene discovery algorithms confronted with blocks of unknown sequence (‘N-islands’) between likely exons are obliged to interpret the N-island as entirely intronic (since otherwise the algorithm would have to posit splice acceptors, donors and exonic sequences entirely composed of ‘N’). If this assignment is correct, then the mature mRNA will seamlessly join exon 1 to the left and exon 2 to the right of the N-island. Bottom: The N-island could harbor an unknown exon X, incorporated into mRNA between exons 1 and 2. **B.** ‘Trinity objects’ are produced by *de novo* assembly of RNAseq reads (REF). Top: if such an object is derived from an exact exon 1-exon 2 splice, then it will align in one continuous high-scoring pair (HSP) in BLASTN analysis to the predicted set of mature transcripts. Bottom: in the case where an N-island contains a hidden exon (Fig. 1A, bottom), the Trinity object assembled from the mRNA should have two HSPs separated by Trinity object sequence; the extra sequence is equivalent to exon X.

We used the Trinity software suite (Grabherr et al., 2011; Haas et al. 2013) to assemble a *de novo* transcriptome for Chlamydomonas, using RNAseq reads that we generated from multiple diurnal cell cycle timecourses (Tulin and Cross, 2015). Since we were primarily interested in evaluating N-islands within gene models, we restricted the RNAseq data input to ~57 million 50 bp single end reads that either (1) aligned to a transcript models that contain at least one N-island, or (2) did not align to any transcript model.

To focus on sequences that are likely to contain useful biological information, we filtered the initial 14,000-sequence Trinity output to retain only sequences that had an e-value (BLASTX) to Volvox or Arabidopsis (TAIR10) peptides of 0.001 or lower were retained. *Volvox* is a recently diverged multicellular relative of *Chlamydomonas,* with approximately 50% neutral sequence divergence but high conservation of many predicted encoded proteins (Prochnik et al, 2010). This reduced the number of sequences to 4412 (median 426 bp; input median 315 bp); many shorter sequences may be either Trinity assembly errors or else too short to generate significant BLASTX alignments. To filter out shorter fragments that were potentially partial versions of longer sequences in the set, we used the uclust software (doi: 10.1093/bioinformatics/btq461). This narrowed the result to a final list of 3114 unique Trinity-generated primary sequences, enriched in longer sequences, and also enriched in potentially conserved protein coding sequence.

These sequences, even after filtering, are likely heterogeneous: some reflect assembly of *in vivo* mRNA species, but there are likely others that are assembly artifacts (for example, track-crossing errors at repeat sequences). We refer to these sequences as ‘Trinity objects’ to reflect initial neutrality concerning their biological status.

To get an estimate of the completeness of the Trinity-generated “transcriptome”, we performed a BLASTN search with Phytozome transcripts against a database of Trinity-generated objects. Since we wanted to resolve intronic N-islands, we used only Phytozome transcripts predicted to splice out N-islands. We will call a splice junction ‘covered’ if a Trinity object aligns with continuous coverage across the junction. Approximately 71% of all positions annotated as splice junctions in the Phytozome reference were covered by a high-scoring pair (‘HSP’), indicating agreement between Phytozome and Trinity objects at those sites that the annotated splicing event occurred *in vivo.*

#### Intronic N-islands: ‘covered’, ‘bridged’ or ‘half-bridged’ by Trinity objects?

In the annotated genome, an intronic N-island is flanked by a splice donor on one side of the N-island, and a splice acceptor on the other side (Figure 1A). This subclass of splice junctions was contiguously ‘covered’ by Trinity objects at only about 60% of the overall the frequency of splice junction coverage in the same transcript set (P≪0.001).

To explain this discrepancy, we hypothesized that some of the 789 intronic N-islands may harbor ‘hidden’ exonic sequence (Figure 1A, bottom). We used BLASTN searches with reference Phytozome transcript models as queries, against a database composed of the Trinity objects. If the N-island contains only intronic sequence, we may find a Trinity object that ‘covers’ the splice junction with a single HSP (Figure 1B, top). This type of BLASTN alignment signals complete agreement between Phytozome and Trinity, and is consistent with location of the entire N-island within a single intron, as indicated by the Phytozome gene model. On the other hand, if the N-island harbors hidden exonic sequence, we may find a Trinity object that ‘bridges’ the splice junction using two HSPs (one on either side of the junction) (Figure 1B). For scoring a bridge, we required two HSPs in the alignment between the Trinity object and the N-island-containing Phytozome transcript aligned to it. We also required extra sequence from the Trinity object between the HSPs. (The second requirement avoids spurious detection of BLAST results with two HSPs that overlap, presumably due to direct repeat sequences at the HSP termini). In addition to the ‘covered’ and ‘bridged’ categories, a ‘half-bridge’ can result if the Trinity object is too short relative to the amount of missing sequence to make a full bridge. We scored each of the 789 intronic N-island splice junctions as either covered (289), bridged (90) half-bridged (9), or ‘no information’ (no Trinity object with coverage relevant to the junction) (398). This result implies that approximately 74% (289/388) of assignments of N-islands as fully intronic in the reference assembly are likely correct, since a continuous Trinity object covers the splice junction containing the N-island (Figure 1B, top). We next turn our attention to the remaining 26%, to determine whether these represent genuine hidden exons, or merely Trinity assembly artifacts.

#### BLAST analysis against *Volvox* to determine evolutionary conservation of bridging sequences

Trinity sequences passing the bridging test above still could represent artifacts of *in silico* assembly. (For example, one could imagine problems from repeats leading to track-crossing. If **R** is a repeat sequence, then an authentic transcript AB**R**CD could plausibly be misassembled with another transcript HIJ**R**KLM**R**OPQ to make the incorrect AB**R**KLM**R**CD, which would score as a ‘bridged’ sequence in the first test above relative to the reference transcript AB**R**CD.)

A sequence from Trinity bridging an N-island should, with some frequency, improve BLAST alignments to the *Volvox* proteome if the sequence contains genuine translated exons. Such alignment improvement should be highly unlikely if the bridge structure is the result of a Trinity assembly artifact. Therefore, we carried out BLASTX with nucleotide sequences corresponding to the aligned HSPs plus the intervening sequence provided by Trinity (‘extended’) as query and the *Volvox* proteome as source of subjects. As a control the intervening sequence was scrambled. We required that queries with and without the intervening sequence both hit the same *Volvox* peptide to avoid situations where an unrelated sequence between the HSPs by chance had a strong *Volvox* alignment. To determine how much score increase we should consider significant, we supposed that small random score changes were equally likely to be slightly better with scrambled controls as with the intact sequence. We observed that score changes were asymmetrical (Supplementary Tables); the scrambled control never showed a score increase > 10, while such increases happened very frequently with the intact sequence. [Note: a BLAST score of 11 is trivially attained in a search of a query against an entire proteome; the reason such a small score is significant in the present case is that the subject for the query is a single *Volvox* peptide pre-selected on the basis of alignment to the flanking regions common to the reference transcript and the Trinity object (Figure 2)]. 69 of 90 sequences that bridge N-islands, and all 9 of the half-bridge N-islands (thus 21% of the intronic N-islands for which we have relevant Trinity information) passed this criterion, and thus are highly likely to contain additional exonic sequence. This criterion is conservative, since a substantial proportion of Chlamydomonas proteins have either absent or patchy alignment to Volvox peptides

**Figure 2.**
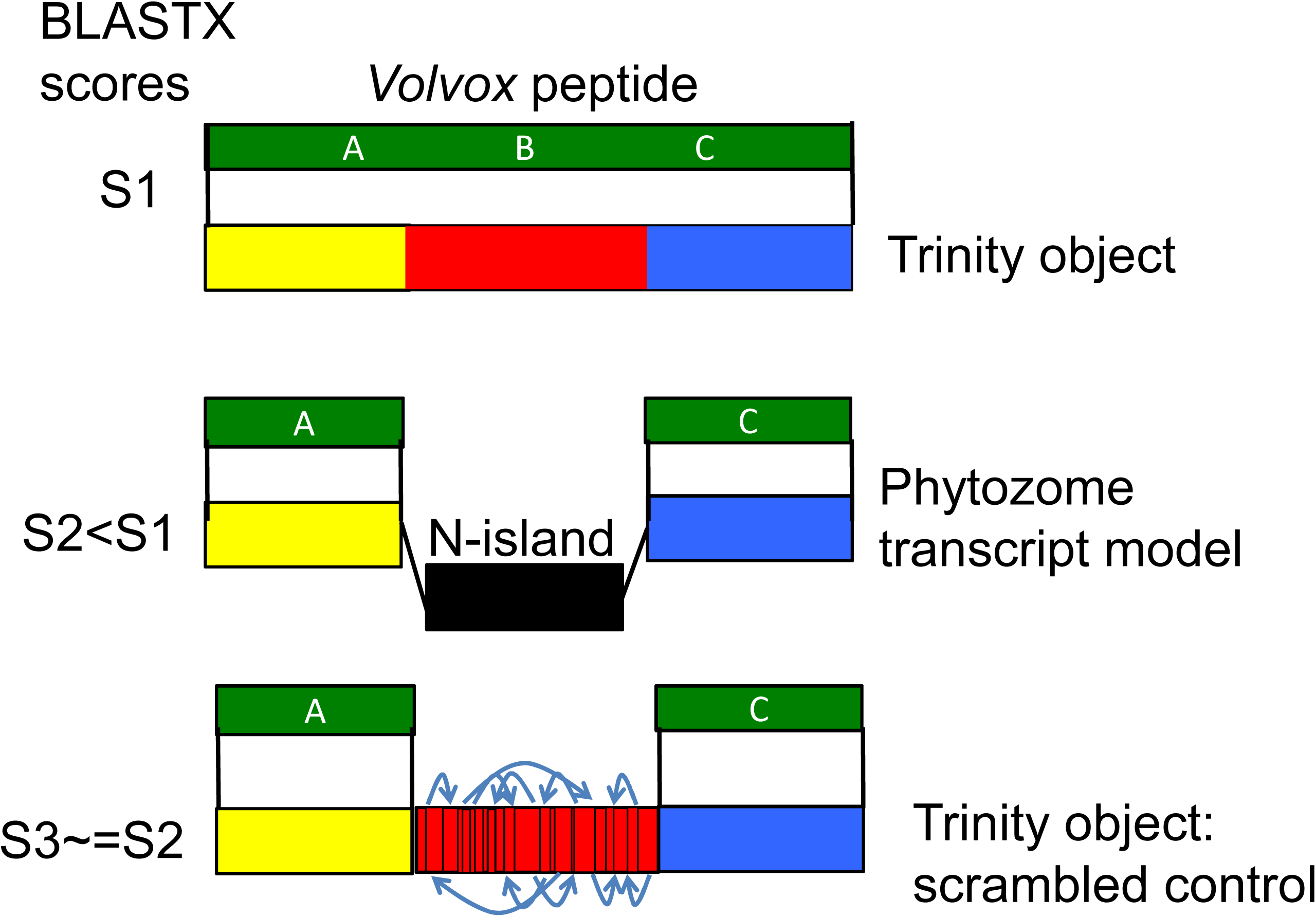
Authentic missing exons can encode evolutionarily conserved sequence: the *Volvox* BLASTP test. Assume that a Trinity object, but not the most closely related Phytozome transcript model, contains a hidden exon (Figure 1). *Volvox* is a close relative to *Chlamydomonas.* If BLASTP of the Trinity object against *Volvox* gives a score S1 due to alignment to segments A,B,C, then BLASTP of the corresponding best-hit Phytozome model (A-C) will give a score S2 < S1, due to lacking alignment from the middle segment lost in the N-island. As a control, scrambling the intervening sequence from the Trinity object (random permutation of the amino acid sequence) should eliminate any scoring advantage, yielding score S3 ~= S2 < S1.

#### Analysis of genomic reads aligned to Trinity objects to determine if candidate missing exons can be explained by RNA splicing

Trinity objects are assembled from short cDNAs copied from cellular RNA. A segment of a Trinity object could have three relationships to genomic DNA: (1) it could be an assembly from RNA sequence transcribed from a single contiguous stretch of DNA; (2) it could be an assembly from RNA transcribed from discontiguous stretches, which are spliced *in vivo* into a single RNA; (3) it could be assembled from RNA transcribed from discontiguous stretches, that are combined by Trinity into a single object by artefactual misassembly *in silico* (most likely by ‘track-crossing’ at repeat sequences, which are extremely abundant in *Chlamydomonas*; see above).

If a segment of a Trinity object was transcribed from a contiguous stretch of genomic DNA, then genomic reads aligned to the segment should tile over a single continuous matching sequence. In contrast, if the segment was transcribed from discontiguous segments, alignment of genomic reads should break into multiple discrete islands. We aligned 100 bp genomic DNA reads to the collection of Trinity objects with Bowtie2 using ‘local’ alignment (which allows alignment even if only a portion of the read aligns to the template). This leads to a highly specific signature of an alignment discontinuity: a ~100-fold excess assignment of ‘starts’ and finishes’ of alignment to nucleotide positions exactly at the point where two discontiguous sequences are joined in the template (Figure 3A,B). These positions were used as seeds to grow a contiguous genomic region out of all reads initially aligned to the Trinity object, by iteratively constructing a majority consensus, discarding reads that fit poorly, and recruiting new reads that aligned well and expanded the consensus. The process terminates when no unrecruited reads both align to the Trinity object and also agree with the growing consensus. The latter criterion is highly effective in discarding reads that align solely by an internal repeat. The result for each discontiguous segment of genomic DNA aligning to part of a Trinity object is an island of sequence supported by a tiled series of reads, with sequence not alignable to Trinity extending from either end (Figure 4A). The extensions are typically of length ~80, equal to the read length 100, minus the minimum amount of sequence required for Bowtie2 to call a local alignment, ~20. Ideally, one such island is found for each discontiguous sequence making up the Trinity object.

**Figure 3.**
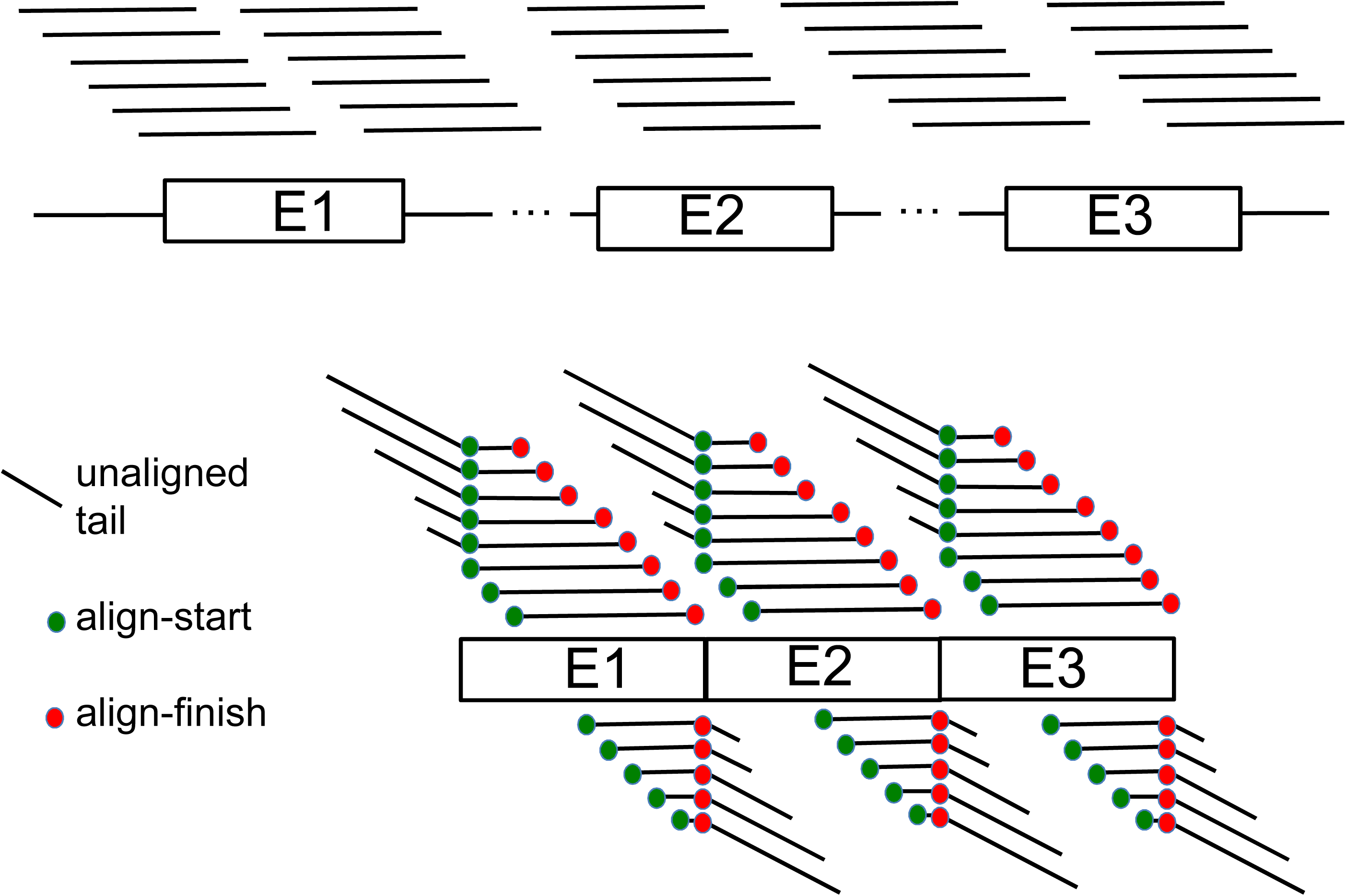
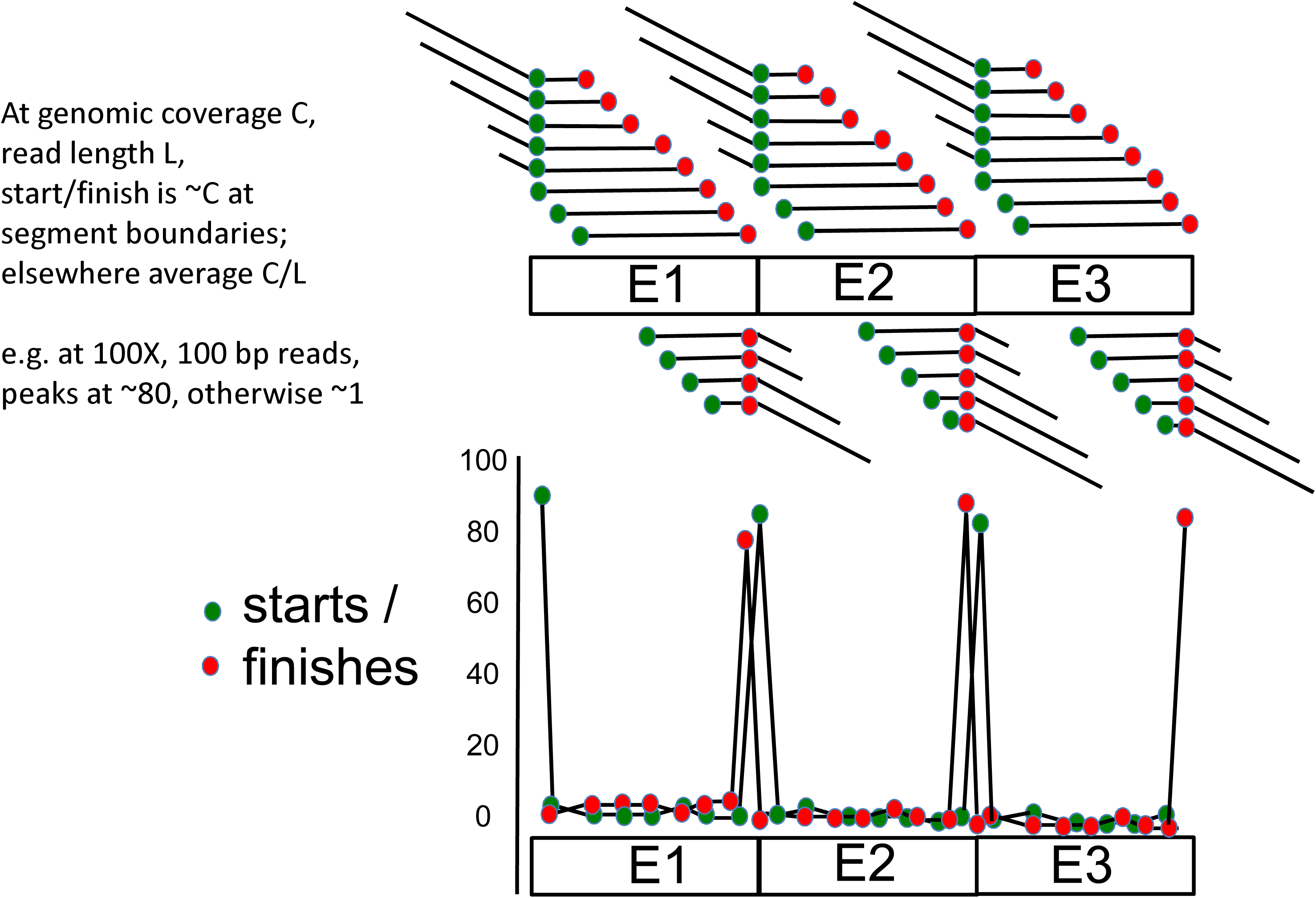
Local alignment of genomic reads to transcripts gives a signature for junctions of discontiguous genomic encoding segments. **A.** Tiled Illumina reads over the genome covering discontiguous segments E1, E2, E3 (top) are aligned locally (not requiring alignment across the complete read) to the fused structure E1-E2-E3 (bottom). These reads have starting and finishing positions of alignment to the E1-E2-E3 structure (green and red dots). **B.** The result for random starts and finishes of reads *in the genome* is a huge enrichment of start positions at the beginnings of the genomically discontiguous segments E1, E2, E3, and for alignment finishes at the ends of E1, E2, E3 (top). This enrichment is on average the read-length (e.g., 100X for 100-bp reads). The plot of starts and finishes of alignments by position then yields a highly sensitive indicator of the segment boundaries.

**Figure 4.**
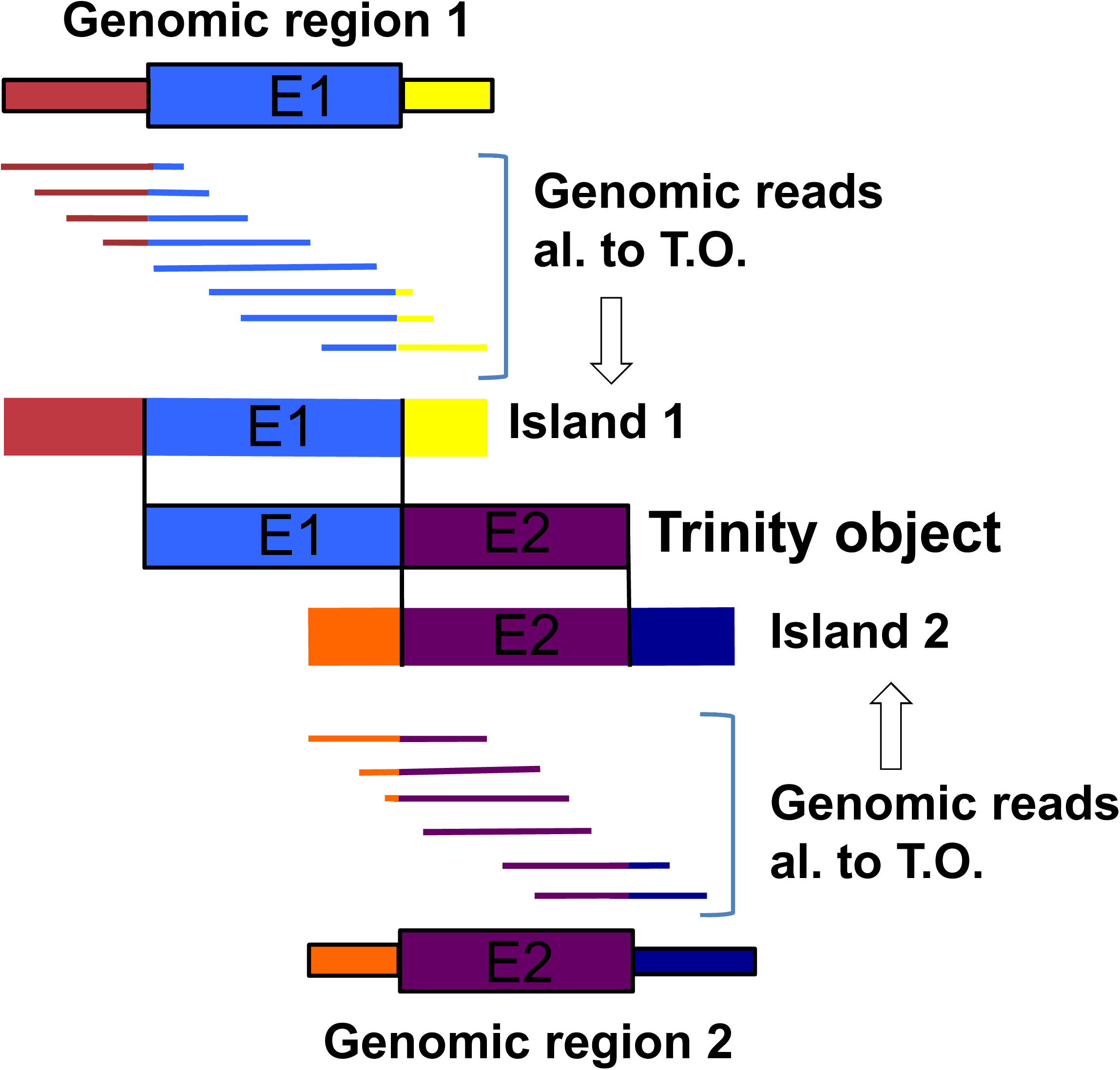
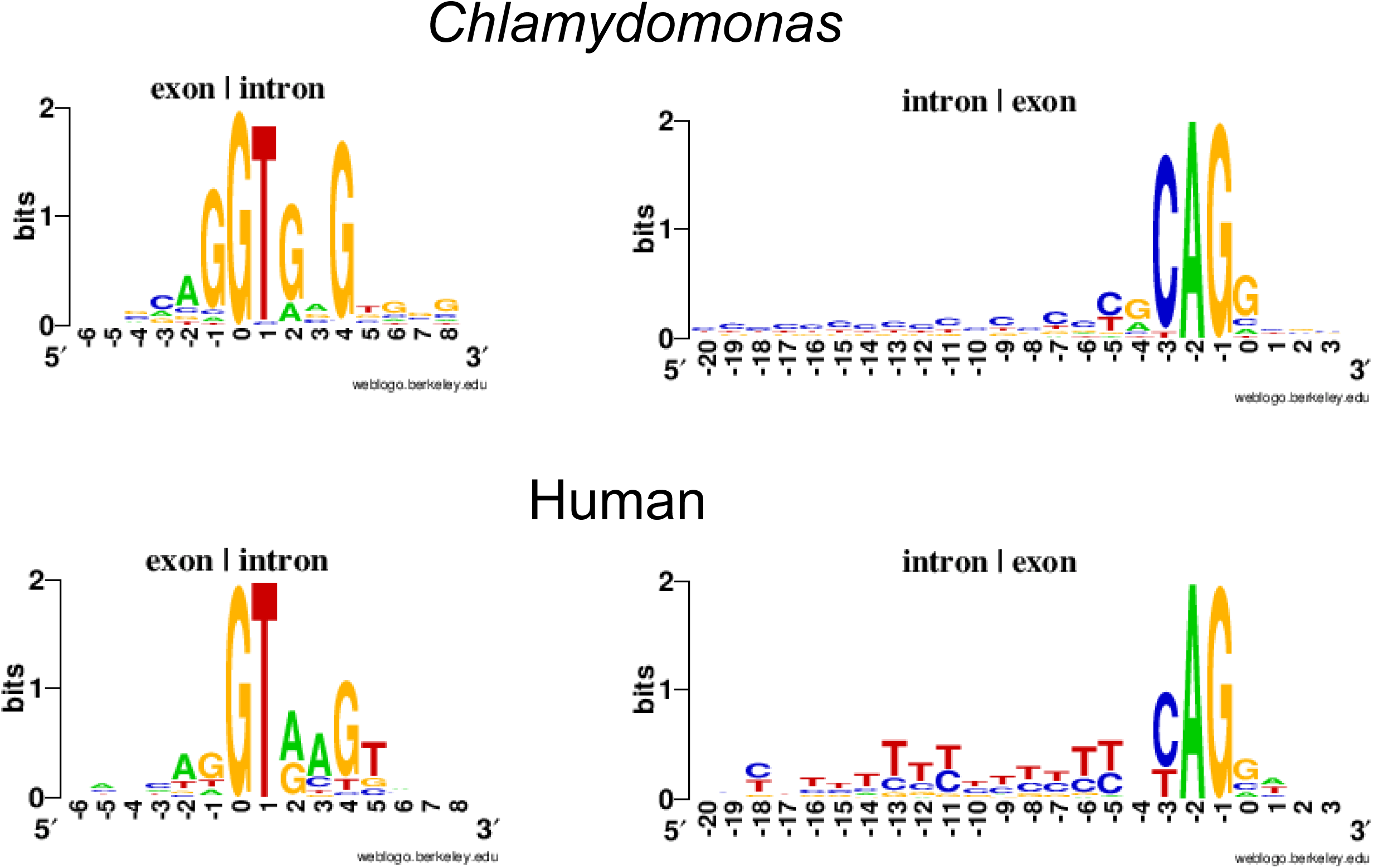
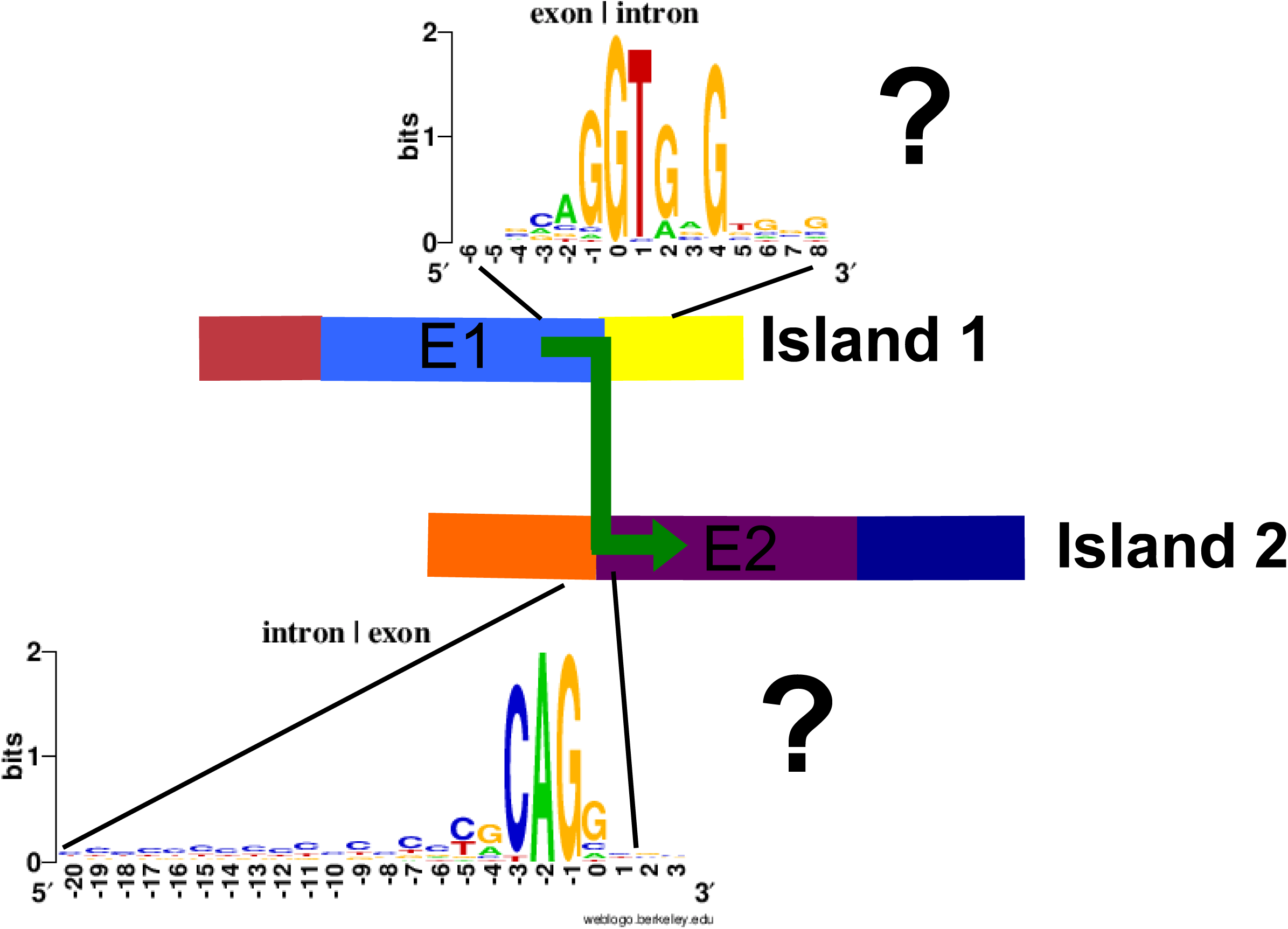
Potential for RNA splicing to assemble consensus islands of genomic sequence into mRNA: the splicing test. **A.** Tiled genomic reads (specific segments indicated by different colors) aligned to a Trinity object encoded by two discontiguous segments E1 and E2 yield island consensus sequences (‘Island 1’, ‘Island 2’). Each island has a central segment identical to E1 or E2 and flanking sequences not found in the Trinity object. Note that when aligned within the Trinity ‘frame’, Islands 1 and 2 have no sequence in common, even though each aligns to a large segment of the Trinity object. **B.** Consensus sequences for splicing in *Chlamydomonas.* Splice sites from the annotated Phytozyme reference genome were collected and analyzed using the Weblogo algorithm (Crooks et al; http://weblogo.berkeley.edu). The human consensus is shown for comparison. **C.** The splicing test. Consensus islands assembled as in Fig. 4B can only be assembled into mRNA by a join at a highly restricted location (in the absence of any terminal direct repeats, a single nucleotide). This provides a highly constrained condition for searching for splice site homology (donor and acceptor). A series of consensus islands connected by high-confidence splices on the same strand forms a ‘connected island’; a candidate hidden exon should be contained within such a connected island to pass the splicing test. Question marks: the test is failed if no match to splicing consensus sequences is detected that can join the islands.

Sequential islands of this kind have no aligned sequence at all comparing them to each other, although each has a central segment aligned to sequential segments of the Trinity object. Alignment discontinuities between these islands (that is, the blue-yellow and orange-purple junctions in Figure 4A) mark an obligatory and exact position where the genomic segments must be joined in order to reconstitute the Trinity object. This restricts positions of potential RNA splicing to one nucleotide. (This is expanded to a few nucleotides in the case of direct repeats at the termini of the Trinity-homologous segments). Internal repeats will typically not yield a neighboring start-finish pair in close enough proximity to explain the join by splicing.

Because the position of the join is highly constrained, we reasoned that if the segments were joined by RNA splicing, then we should be able to detect sequence signatures permissive for splicing specifically at these junctions. To determine this signature, we examined the genomic splice site sequences in the reference assembly/annotation to determine a splicing consensus (Figure 4B,C). Consistent with previous work in many eukaryotic organisms, the strongest determinant was the /GT&.AG/ dinucleotide pair at intron termini; other flanking sequences also contributed significantly to the consensus, equivalent to almost 8 nucleotides of specified sequence. We found that a match score of 11 bits (obtained by summing information content for observed nucleotide at each position) was attained by ~99% of Phytozome annotated splice junctions, and by only ~1% of randomly selected genomic sequences, and we chose this value as a cutoff for splice junction detection. Because of the near-absolute requirement for the /GT.(intron).AG/ dinucleotides for splicing, we imposed a two-step test: first, presence of the dinucleotides; second, a score of at least 11 (the dinucleotides alone give a score of 8). We applied this test to the discontinuities in alignment of genomic sequences to Trinity objects, as detailed above.

The results of this procedure for four typical cases with a ‘bridged’ N-island (Figure 1B, bottom) are shown in Figure 5A. In these diagrams, the successive red bars at top represent islands aligned to Trinity; the black marks at left and right of these islands represent non-Trinity-homologous sequence, which in general is flanking intronic sequence. The blue arrows are high-confidence splice junctions according to the test described above. In the four typical cases in Figure 5A, the algorithm has assembled multiple candidate exons. In all cases these exons could be joined by high-confidence splices, thus these sets of sequences form ‘connected islands’ that could be assembled into a single mature transcript by splicing.

**Figure 5.**
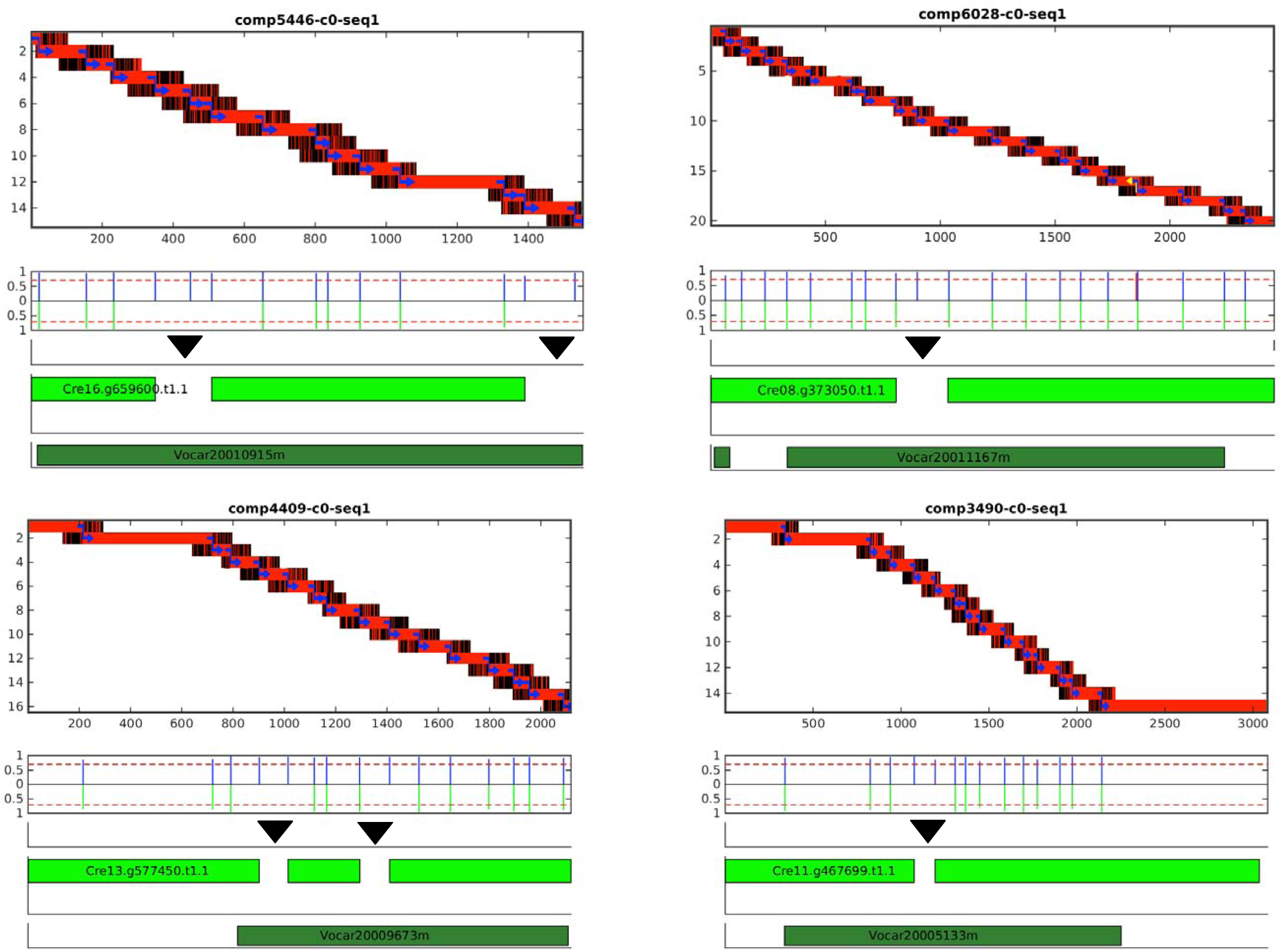
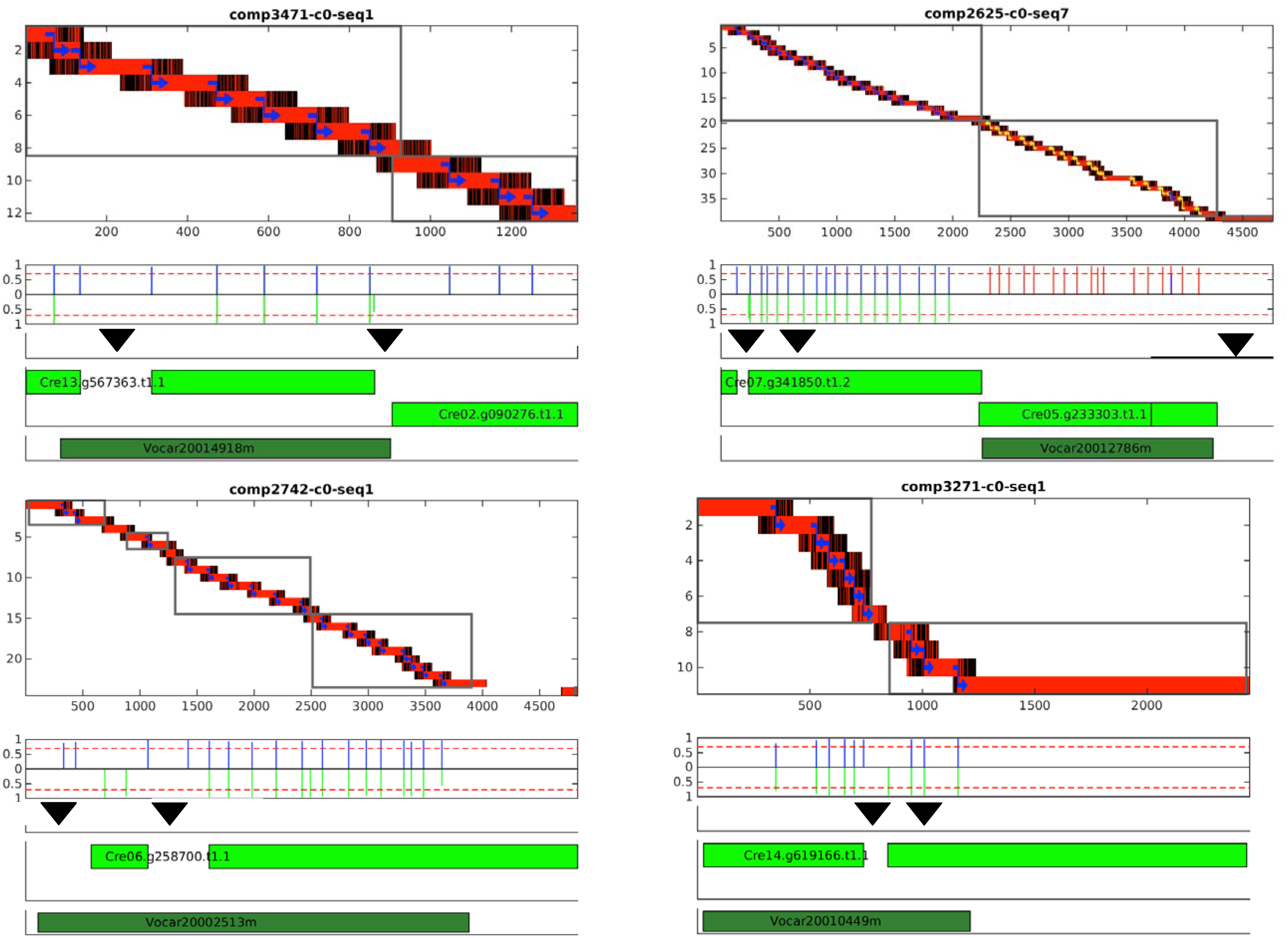
Graphical summary of analysis of Trinity objects. The ‘frame’ for all data is the sequence of the Trinity object (numbers on x-axis of top graph). At top is the result of analysis of genomic reads aligned to the Trinity object. The red/black bars are alignment islands (Figure 4A), where red indicates identical sequence to Trinity and black non-identical sequence. The blue joining arrows indicate high-scoring splice junctions (if detected, junctions on the opposite strand are in yellow). Below is a quantitative trace of splice junction locations and scores (standardized to the maximum possible score of ~15; the cutoff value for acceptance of 11 is illustrated by the red line). The blue lines are splices in our model; green lines below are splices from the reference transcript model aligned to the Trinity object. The green bars indicate regions of alignment (HSPs) to the specified Phytozome reference transcript. The black triangle indicates the approximate location of putatively spliced-out N-islands in the reference. The bottom graph indicates the positions of alignment to the *Volvox* proteome. **A:** typical cases for ‘bridging’ Trinity objects. In these four cases the entire set of consensus islands are joined into one ‘connected island’ by same-sense splices. **B:** rare aberrant cases where the Trinity object is due to ‘track-crossing’ *in silico,* and contains regions of two distinct reference transcripts (top two examples), or where splice detection fails even on the same reference transcript (bottom two examples). In these cases there are multiple connected islands (grey outlines).

Also indicated in the figures are positions of splice sites in the aligned reference annotation compared to those detected in the island termini. The splice junctions are almost always are on the same strand; one case where a splice junction could be called on both strands can be noted in Figure 5A, top right (yellow arrow).

The exact lineup of almost all computed and reference splice junctions constitutes strong evidence for accuracy of our procedure, as well as supporting all these intron locations in the reference annotation. The exceptions to this rule are essentially all detected at the position of N-islands thought to be spliced out according to the reference, that are in ~25% of cases associated with additional exons in this analysis.

Examples of the main types of occasional aberrant patterns are shown in Figure 5B.) The boxes delineate ‘connected islands’ defined by splice junctions on the same strand. Top left and top right: evident Trinity assembly errors in which two transcripts were fused computationally into single Trinity objects. Note that in the case at top right, the fusion switched the strand of the sequence, as indicated by splice junction orientation (blue to yellow). The bottom two cases are more likely to be simple failure of detection of splice junctions by the algorithm.

#### Case studies: *PF20* and *CDC27.*

We will discuss two cases of exon discovery in detail. *PF20* (required for flagellar function) was chosen as an example since the complete cDNA sequence encoding 606 amino acids was described previously (Smith and Lefevbre (1997)), but we noticed an intronic N-island in the annotated reference genome (between exon 8 and 9 in the Phytozome gene model Cre04.g227900) in the annotated reference genome that apparently eliminated coding sequence for amino acids 357 to 488.

Figure 6A diagrams our results for *PF20,* encoded as described above for Figure 5. Our results replace the N-island with 4 missing exons. Inclusion of these exons in the predicted mRNA results in a complete coding sequence exactly matching the results of Smith and Lefevbre (1997) (Figure 6B). The bottom dark green bar in Figure 5A indicates regions of homology to *Volvox* PF20. It is clear that the bridging sequence is required for optimal alignment to *Volvox.* It is important to note that this result, recapitulating a careful single-gene study, was generated by unsupervised computations, with the only input being Illumina libraries from bulk cDNA and genomic DNA.

**Figure 6.**
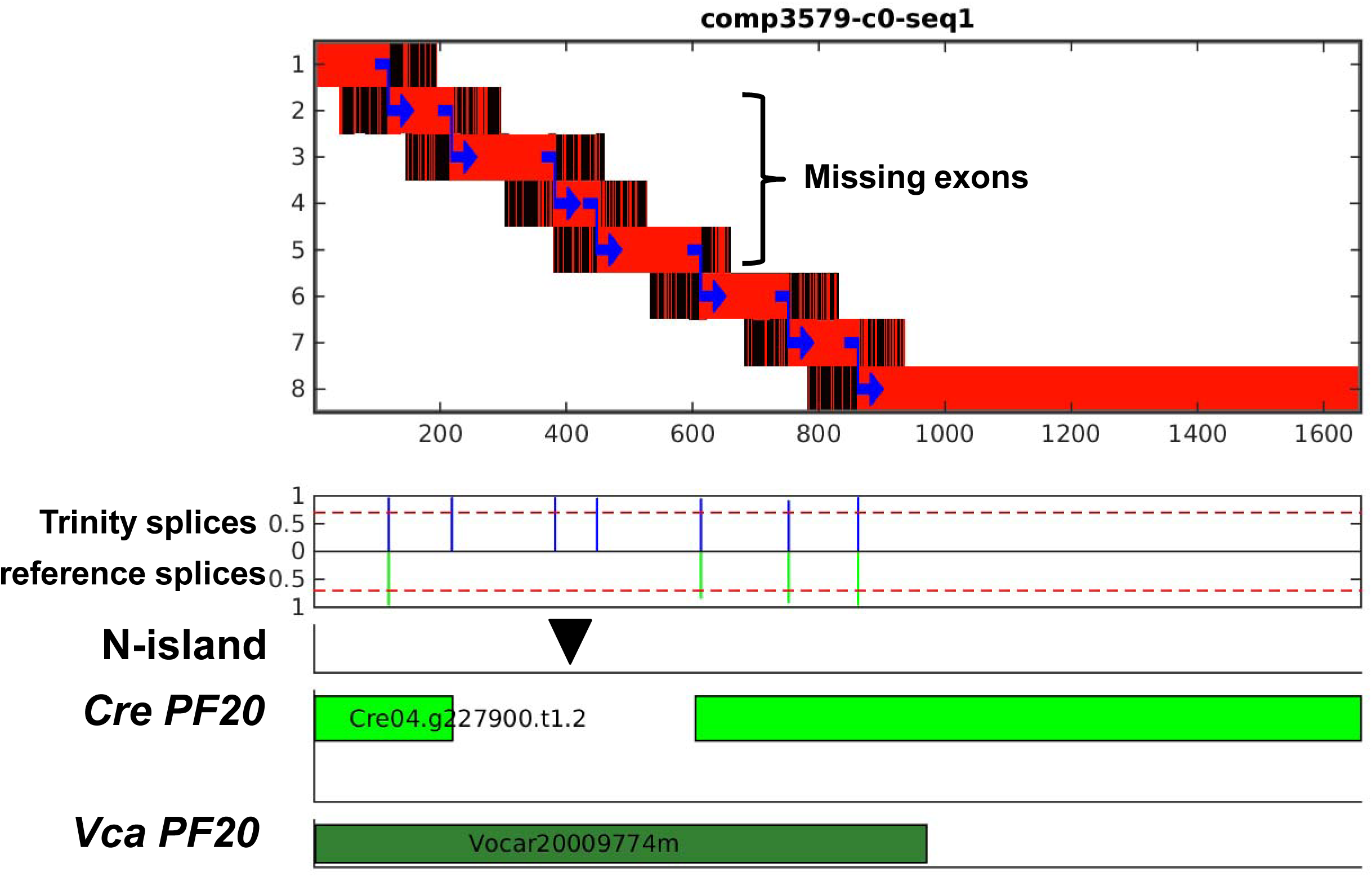
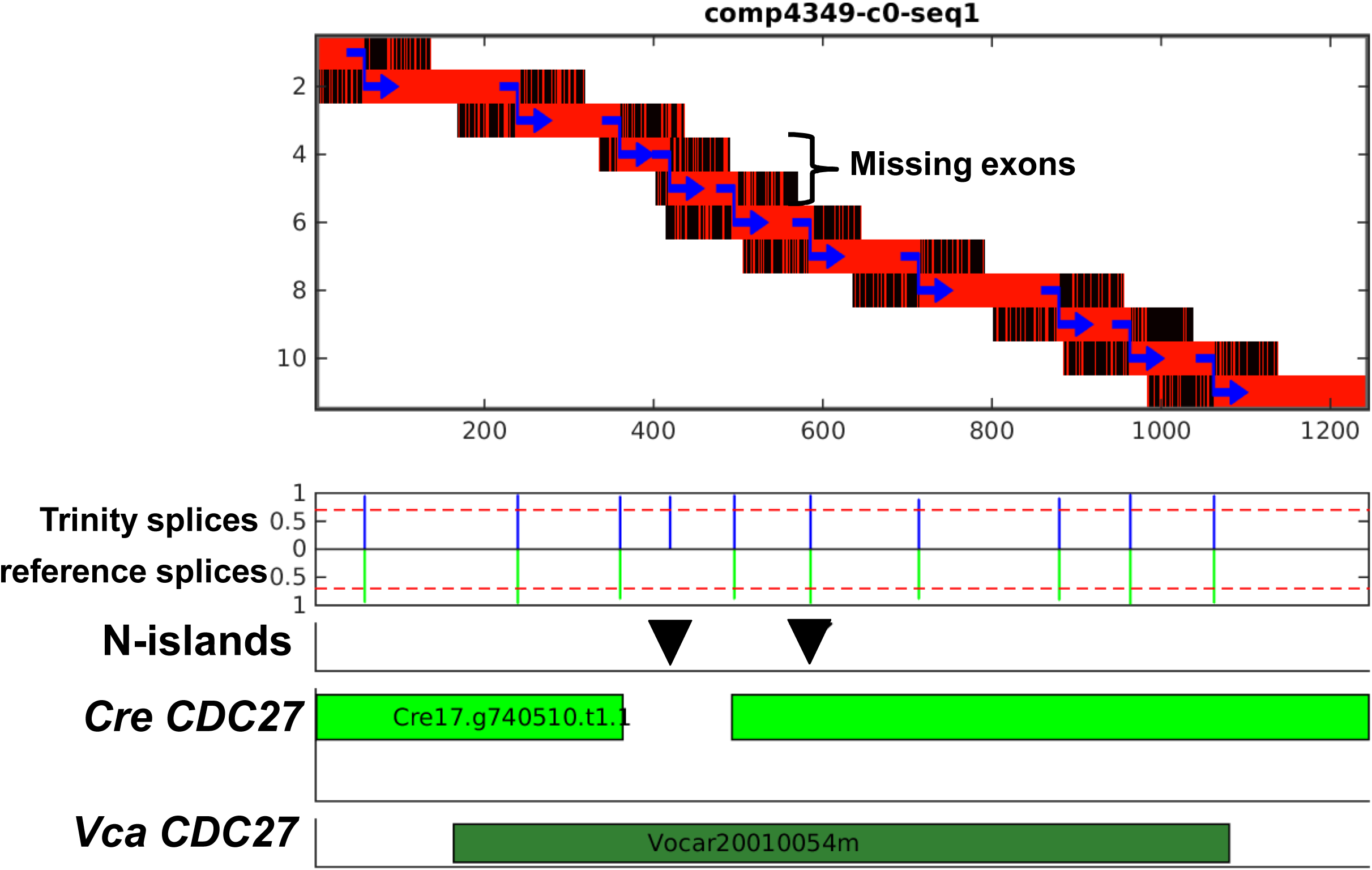
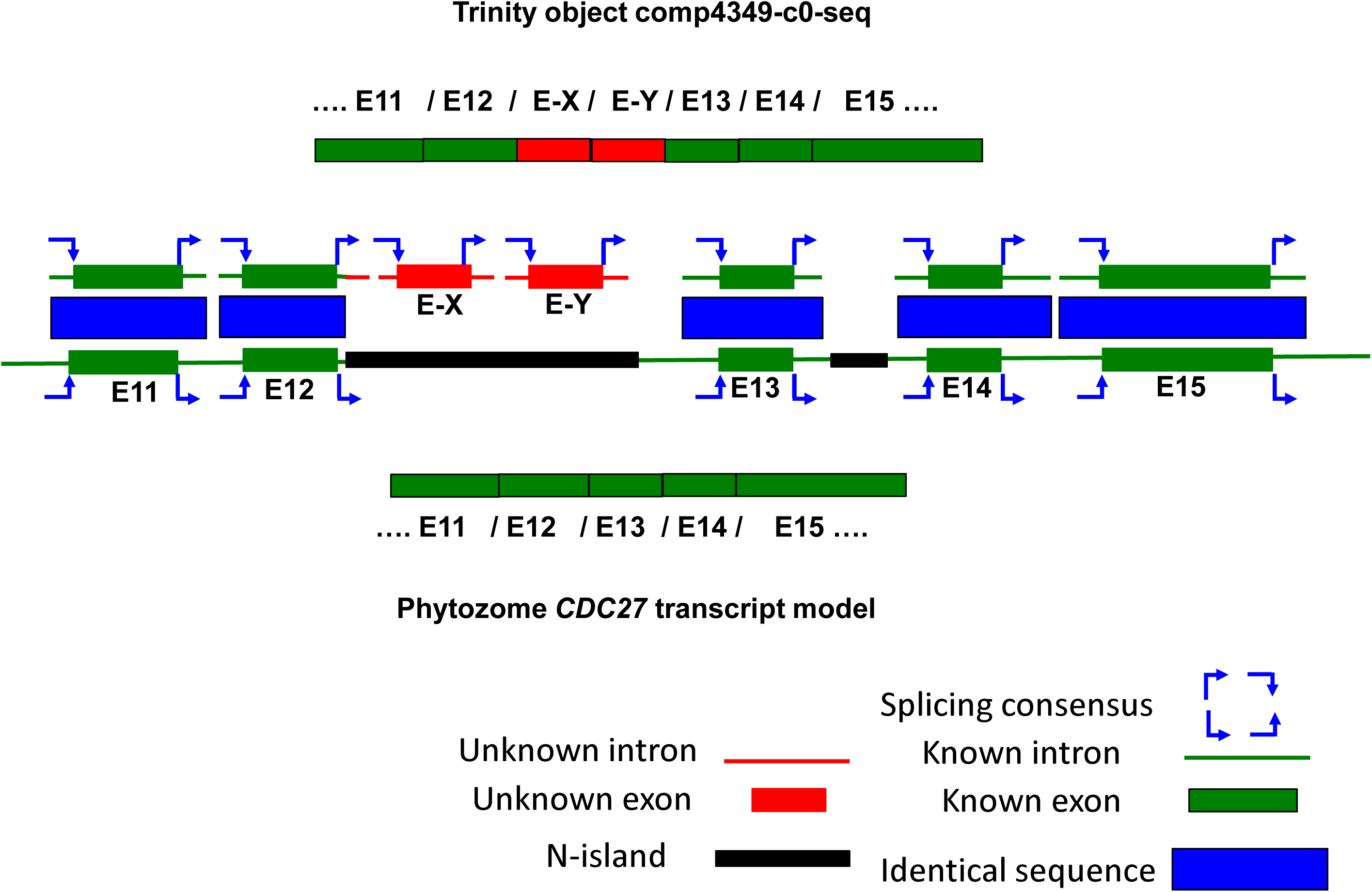
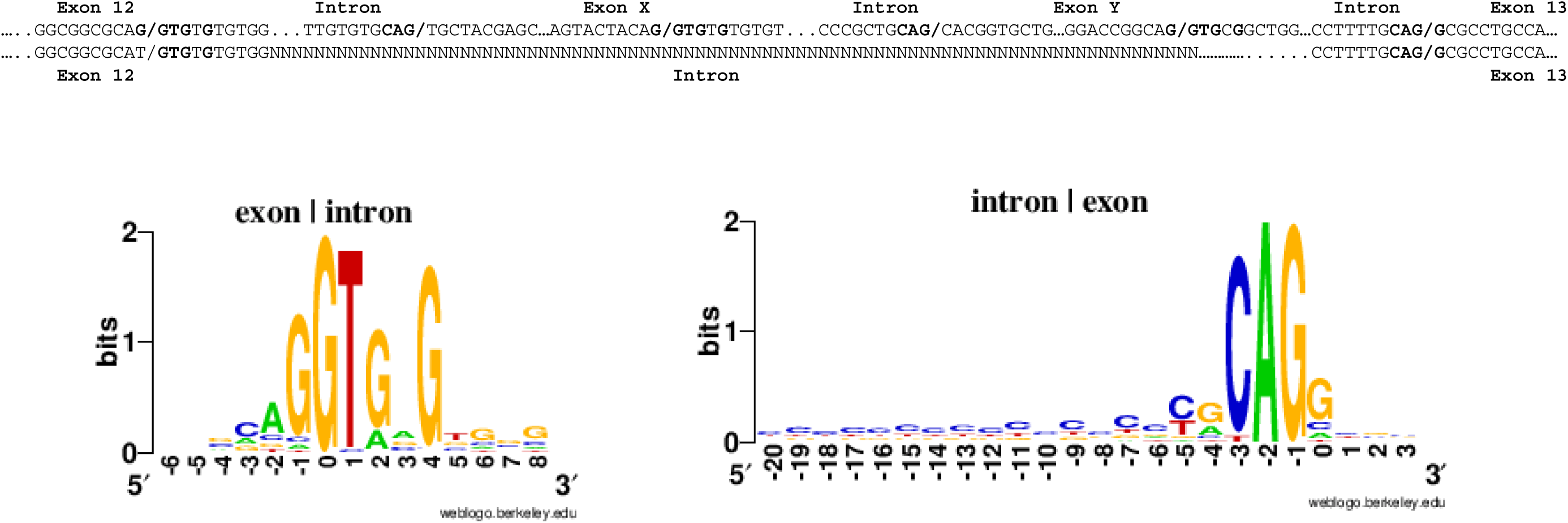
N-islands can contain hidden exons. Two examples are shown: **A:** *PF20;* **B,C:** *CDC27.* **A,C:** diagrams as in Figure 5. In both cases, one N-island contains hidden exons (four in *PF20,* two in *CDC27*). The second N-island in *CDC27* is covered; the diagram shows it to be authentically intronic according to the Trinity object analysis. **B**. A portion the translation of the Phytozome model for *PF20,* the PF20 amino acid sequence determined from cDNA by (Smith and Lefevbre 1997; http://www.uniprot.org/uniprot/P93107), and the translation of the Trinity object. A black bar indicates the position of the intronic N-island in the Phytozome gene model. **D.** The analysis recovers splice-site-proximal intronic sequences; illustrated for *CDC27.* The islands from genomic reads, including Trinity non-homologous sequence, align perfectly with reference genomic sequence for individual exons (green bars) and surrounding intronic sequence (green lines). Similar extensions (red lines) beyond the Trinity-aligned segments (red bars) are detected for the two hidden exons ‘E-X’ and ‘E-Y’. **E:** these extensions are characterized by strong terminal splice site homology, which we take as evidence that ‘exons X and Y’ are in fact spliced into the mature *CDC27* mRNA as reflected in the Trinity object comp4349-c0-seq1.

For a second example, CDC27 (*Chlamydomonas* Cre17.g740510) is a conserved component of the anaphase-promoting complex (Zachariae and Nasmyth 1999). The Cre17.g740510 gene model has two intronic N-islands, between exons 11 and 12 (246 N) and between exons 12 and 13 (100 N). We identified a Trinity object which covered exons 9 to 19 of the CDC27 transcript model (Figure 6C). This alignment had two separated HSPs surrounding the first N-island, with extra sequence between the HSPs in the Trinity object (the signature for possible hidden exons [Figure 1]). The first HSP aligns to reference exons 9-11 and the second HSP covers reference exons 12-17 (with all splice junctions) including the 3 prime UTR. The intervening “missing sequence” between exons 11 and 12 is accounted for by splicing in two new exons ‘X’ and ‘Y’ from a previously unknown genomic segment; the extra peptide sequence thereby added to CDC27 is supported by *Volvox* homology (Figure 6C). Indeed, this extra peptide sequence is conserved in CDC27 from a wide range of eukaryotic species including *Arabidopsis,* animals and yeast. In contrast, the second annotated intronic N-island (triangle in Figure 6C) is ‘covered’ – our analysis is in agreement that this N-island is completely contained within an intron.

We noted above that the Trinity-non-homologous ‘tails’ at 5’ and 3’ ends of the aligned islands are likely to be intronic. Indeed, these sequences for ‘covered’ introns exactly matched reference splice donor and acceptor and neighboring intronic sequences at each covered splice junction (Figure 6D,E) – thus, splice junction-proximal intronic sequences are accurately recovered by local alignment of genomic reads to Trinity objects, independent of any prior reference or annotation.

#### Exonic N-islands

The large majority of N-islands are annotated as intronic. There are, however, 40 exonic N-islands in the reference assembly. Most are assigned to 3’UTR or 5’UTR. We found four cases where a Trinity object bridged an exonic N-island; in three cases this bridge was in coding sequence, and resulted in a significant improvement in BLAST score against *Volvox.*

#### Flanking N-islands

The intronic and exonic N-islands examined above were all annotated within the boundaries of a gene. There are N-islands in the annotation that flank an annotated gene; it is clearly possible that a gene might actually extend into (or even beyond) a large region of unknown sequence immediately to its left or right. We defined a flanking N-island as an N-island starting within 100 bp of the 5’ or 3’ borders of a gene model. If the authentic gene extends into the N-island, then there could be a ‘half-bridging’ Trinity object aligning at one end to the reference gene model, with the other end ‘hanging’ in the N-island. We identified 181 flanking N-islands in the genome, 11 of which could be composed as a connected island extending into the N-island, significantly increasing the *Volvox* BLAST score. Two examples are presented in Figure 7. Because the Trinity objects are short, it is unlikely that these extensions complete the transcript model, and the lack of a full bridge intrinsically makes these hits less reliable, but it is clear that extension of these methods could significantly extend or even complete gene models by foraging into the adjacent N-islands. We suspect that the low yield of these structures is related to the sequencing difficulties (simple sequence, repeat regions) that led to them being called as flanking N-islands in the original annotation.

**Figure 7.**
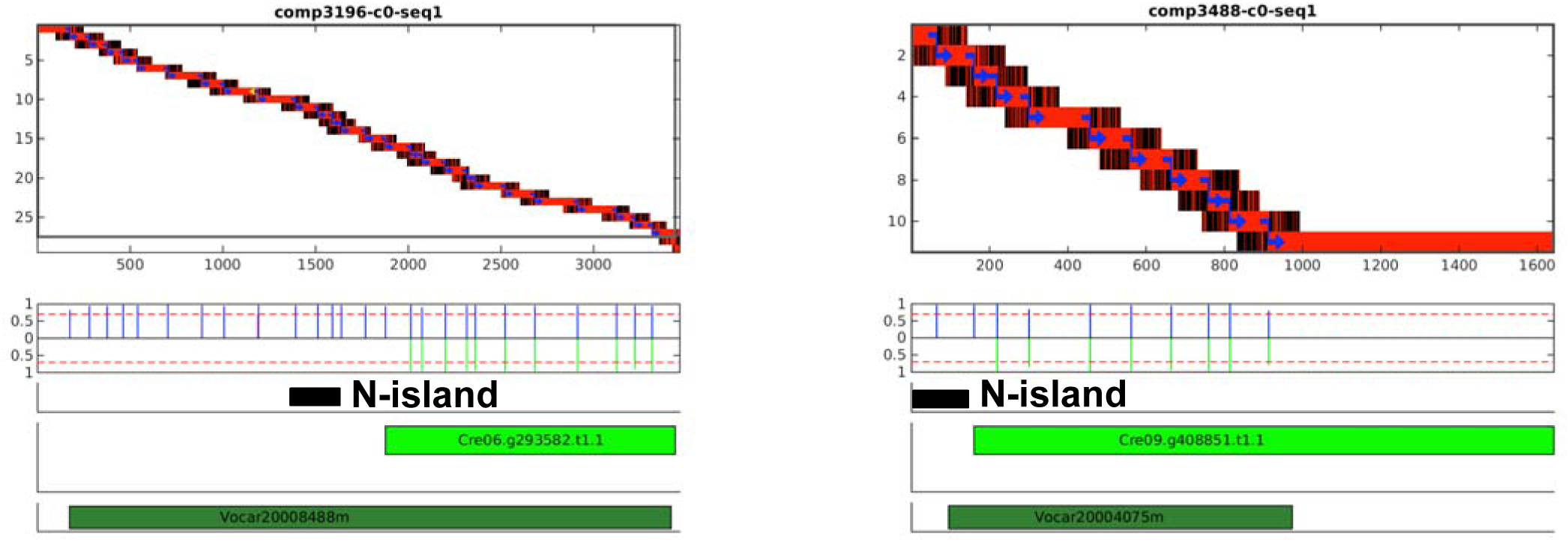

## DISCUSSION

Our analysis suggests that a substantial majority of assignments of unknown blocks of sequence to intronic regions are in fact correct. In some cases, though, our results suggest additional internal coding exons, or N- or C-terminal extension of coding sequence, to replace the putatively non-coding N-islands.

The methods used here are useful for ‘proofreading’ a reference assembly/annotations, but the methods do not require a prior reference. In principle, these methods could validate RNAseq-based transcriptome assemblies, including providing genomic splice sites and flanking intronic sequences, even for organisms without any assembled genome or annotation.

In practice, while our computations per se did not use the assembled genome or reference annotation, the results could be evaluated with much higher confidence when looking for a revision of the current assembly, rather than entirely *de novo* sequence. Thus, we have more confidence in the bridged than the half-bridged cases (such as with the flanking N-islands), and we did not so far attempt to evaluate Trinity objects that were completely unalignable to any reference transcript. One limitation to our approach was the use of single-end, non-strand-specific 50 nucleotide RNAseq reads. We expect that longer, paired-end and perhaps strand-specific libraries might improve resolution and reduce track-crossing sufficiently that a reliable full transcriptome, along with junctional sequences at promoter, termination and splice sites, might be recovered *de novo.* The *Chlamydomonas* genome is a favorable case in that it is 1.2 *10^^^8 base pairs, of which ~one-third is exonic. It is unfavorable in having a very high density of repeats and low-complexity sequence (likely the source of most alignment difficulties including N-islands, and certainly the source of most difficulties we encountered with our approach).

In principle, the genomic islands recovered to account for Trinity objects could themselves be used to align genomic reads, ultimately finding the next exon and providing sequence of the entire intron. We have not attempted this because of the clear danger of track-crossing in repeat sequences, which are especially enriched in intronic sequence.

The test for splice sites at the borders of the Trinity-nucleated genomic islands tests the validity of the Trinity object, since it appears to be a sensitive indicator of track-crossing leading to fusion transcripts (Figure 5B). This could be a useful proofreading step when Trinity-generated transcriptomes are used in the absence of a reference genome sequence, since it requires only a single library of genomic reads aligned to the Trinity objects, and requires no prior assembly of a reference genome.

The combination of *de novo* transcriptome assembly followed by alignment of short genomic reads onto the assembled transcriptome objects in principle represents a massively parallel approach to validating a genome assembly/annotation, equivalent in some ways to checking a near-complete collection of ESTs against the annotation in a single experiment. Our implementation was not quite up to this challenge, because the assembly was only ~70% complete, and a significant minority of the assembled objects had track-crossing and other issues (Figure 5B). However, these seem likely to be solvable issues, through the use of better RNAseq libraries, and through making use of paired-end information for both RNAseq and DNA libraries.

While our approach did uncover some novel exons missed in the prior assembly/annotation, overall our results document the very high quality of the annotation. Most N-islands annotated as intronic really are purely intronic, and almost all annotated splice junctions are confirmed: they are covered by continuous Trinity object sequence, and put together from discontiguous genomic segments with high-confidence splices.

The current assembly/annotation reports ~4 Mb of N’s (3.5% of the genome), distributed in 1,343 islands of highly variable size. 63% of these islands are located in introns, and these are primarily the N-islands the current study addresses. However, these intronic N-islands are short (only 15% > 1 kb, none > 10 kb) account for only 14% of the total N’s; most N’s are found in large islands (40% > 1 kb, 32%>10 kb) annotated as intergenic. It is likely that at least some of these large unknown regions contain unknown transcripts. Such transcripts should be represented in our study by Trinity objects that did not align to any current Phytozome transcript model. We have not examined this question in detail, but it is clear that few such objects exist in our dataset, potentially accounting for no more than a very small proportion of the 3.7 Mb of N’s in intergenic islands. Therefore, while we cannot make a strong conclusion on this point, it seems at least plausible that most of the large intergenic N-islands are composed of non-coding sequence; this is a reasonable result, since low-complexity or repeat-containing sequence is both relatively unlikely to contribute to stable coding mRNA, and also is likely to cause the sequencing/assembly difficulties that generate large N-islands.

## Acknowledgments

Funding support was from PHS 5RO1-GM078153.

